# Endothelial CLK2 as a therapeutic target for acute radiation syndrome

**DOI:** 10.1101/2025.11.05.686891

**Authors:** Ryan R. Posey, Alican Özkan, Yuncheng Man, Jessica F. Feitor, Amanda Jiang, Jie Ji, Nina Theresa LoGrande, Christos Kyprianou, Anna M. Howley, Bogdan Budnik, Jonathan D. Lee, David B. Chou, Donald E. Ingber

## Abstract

Damage to the vascular endothelium is a major contributor to acute radiation injury in multiple organs that underlies acute radiation syndrome (ARS), yet there are no FDA-approved radiation countermeasures targeting endothelial cells. Use of kinome-scale CRISPR screens performed in cultured human vascular endothelial cells isolated from different organs identified CLK2 as a potential radioprotective target. Pharmacological inhibition of CLK2 using TG003 and Cirtuvivint protected these endothelial cells against radiation injury and reversed its effects across the transcriptome and phospho-proteome. Human Organ Chip models of human intestine and lung that contain organ-specific epithelium and microvascular endothelium faithfully replicated clinical features of ARS when exposed to radiation, which were prevented when treated with CLK2 inhibitors. Thus, CLK2 inhibitors may represent a new class of radiation countermeasure drugs that can protect multiple organs against radiation-induced toxicities in patients with ARS.

---

Acute radiation syndrome (ARS) encompasses a myriad of radiation injuries affecting the gastrointestinal, pulmonary, hematopoietic, neurovascular, and integumentary systems [1, 2], with organ-specific toxicities being observed at different doses of radiation exposure in different organs. The hematopoietic and gastrointestinal systems exhibit the greatest sensitivities, with full effects observed starting at doses of ∼1 and ∼8 gray (Gy), respectively [1], with the lung displaying injury responses at a high dose exposure (>8-10 Gy)[3]. Unfortunately, only four therapies are currently approved by the FDA for ARS, all of which target the hematopoietic system: Neupogen (G-CSF), Neulasta (PEGylated G-CSF), Leukine (GM-CSF), and Nplate (TPO-receptor agonist) [4–6]. Mechanistically, these therapies have been selected for their ability to combat toxicity due to cytopenias observed in ARS. However, many cell types and tissues that are highly susceptible to radiation injury, such as gastrointestinal and pulmonary epithelia and endothelia, lack therapies developed specifically to promote their survival and protect tissue functioning. Thus, the development of therapies that target these cell types represents an untapped reservoir of possible medical radiation countermeasures.

The vascular endothelium has been shown to mediate acute radiation injury and inflammation, particularly in the gastrointestinal tract, with the microvasculature being more radiosensitive than the parenchymal cells[7–10]. Recent studies with human organ-on-a-chip (Organ Chip) microfluidic culture models have confirmed the key role of the endothelium, which significantly enhanced sensitivity of adjacent epithelial cells to radiation injury in human intestine [10] and lung [11], corroborating observations from previous *in vivo* studies in animal models [12, 13]. Therefore, protection of the endothelium from radiation injury could represent a promising therapeutic avenue for treatment of ARS in multiple organs. However, numerous mechanisms by which the endothelium can alter radiation sensitivity of neighboring parenchymal tissues have been proposed, including activation of inflammatory leukocytes via cytokine production [14, 15], adhesion molecule upregulation [16–18], increased barrier permeability [19, 20], induction of coagulopathies [21, 22], dysregulation of metabolic pathways [23], increased reactive oxygen species (ROS) production [24], altered calcium homeostasis [25], and ischemia [26]. Given this multitude of possibly injurious mechanisms, it is difficult to know where to begin when seeking a therapeutic target. Therefore, we pursued a mechanism-agnostic, but endothelial cell-targeted, approach using a kinome-scale CRISPR screen, rather than focusing on the transduction mechanisms themselves.

## Kinome-Scale CRISPR Screens Identify Potential Novel Radiation Countermeasures

To identify possible therapeutic targets in primary human endothelial cells, we use CRISPR/Cas9 to target the human kinome, which is composed of kinases that can be modulated by at least 243 kinase inhibitors previously described in the clinical literature [27]. Pooled screens with CRISPR/Cas9 enable high-throughput dissection of genotype-phenotype associations; however, they are most commonly performed in immortalized cell lines [28–30] because primary cell screens are often limited by the quantity of cells and poor editing efficiency. To overcome this limitation, we cloned the mGreenLantern protein in place of GFP in the LentiCRISPRv2-GFP vector (LCv2-GL) (**Supplementary Fig. S1A**), which provided superior signal-to-noise compared to LentiCRISPRv2-GFP (**Supplementary Fig. S1B**). This resulted in transduction with high efficiency in three different types of primary endothelial cells — human small intestine microvascular endothelial cells (HSIMEC), human pulmonary microvascular endothelial cells (HPMEC), and human umbilical vein endothelial cells (HUVEC) (**Supplementary Fig. S2**). The ability of this vector to induce a gene knockout in primary human endothelial cells was confirmed by transducing HSIMEC with lentiviruses containing LCv2-GL with an sgRNA against *TP53*, which resulted in a nearly complete knockout 7 days post-transduction, as detected by Tracking of Indels by Decomposition (TIDE) (**Supplementary Fig. S3**).

We then cloned 3152 sgRNAs targeting 763 kinases, with 4 guides per gene and 100 non-targeting guide sequences into LCv2-GL (**Supplementary Fig. S4** and **Supplementary Table 1**). To determine the parameters of our screen, a range of γ-radiation doses were tested for their effects on human endothelial cell survival. To be as stringent as possible while allowing sufficient cell coverage after the screen, a radiation dose of 4 Gy was chosen as ∼15% of two different types of endothelial cells (HSIMEC and HUVEC) survived at this dose (**Fig. 1A**). For HPMEC, 16 Gy was selected based on our previous work showing this to be the optimal radiation dose to produce endothelial injury in human Lung Chips lined by HPMEC [31]. Using the MAGeCK Flute pipeline for normalization and quality control (**Supplementary Fig. S5**), each type of endothelial cell was treated as an independent condition in our design matrix (**Supplementary Table 2**). Importantly, we identified highly overlapping gene-level hits that were shared by the different endothelial tissue types (**Fig. 1C**). We also observed a strong correlation between effect sizes and statistical significance across all endothelial subsets tested (**Supplementary Fig. S6** and **Supplementary Table 3**).

**Figure 1.**
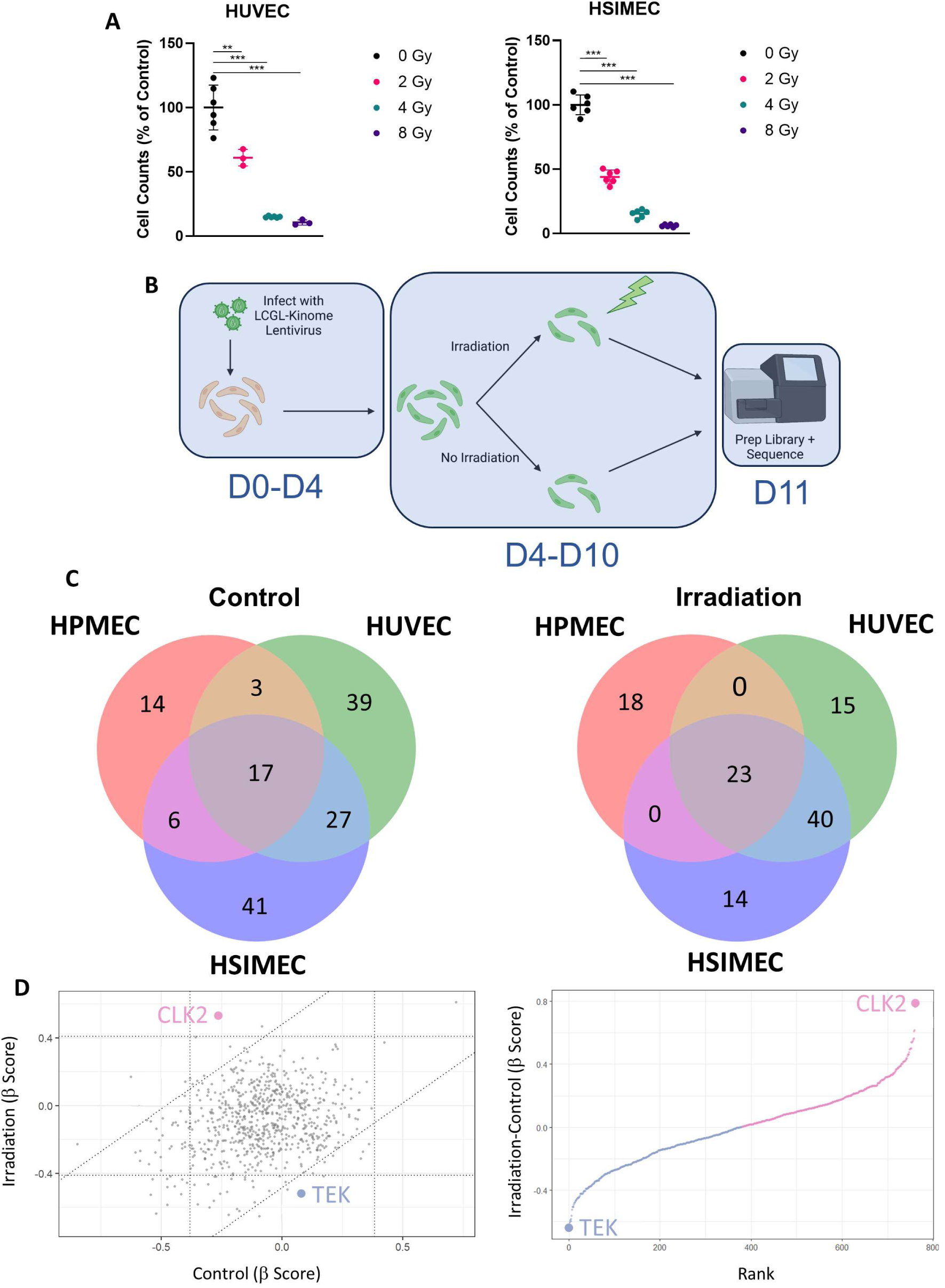
Design and optimization of a kinome-scale radiation resistance screen in human endothelial cells. (**A**) Radiation dose titration in human umbilical vein endothelial cells (HUVECs) and human small intestinal microvascular endothelial cells (HSIMECs) showing effects on cell number (Cell Counts) in response to exposure to 0, 2, 4, or 8 Gy radiation (n = 6;**, p < 0.01; ***, p < 0.001). (**B**) Schematic overview of the CRISPR Screen workflow and timeline. (**C)** The number of statistically significant sgRNA targets identified in each screen under control and irradiated conditions in HUVECs, HSIMECs, and human pulmonary microvascular endothelial cells (HPMECs). (**D**) 9-square plot showing top hits from pooled screen analysis (left) and gene-level ranking (right) from MaGECK Flute.

The Angiopoietin-1 (Ang-1) pathway has been shown to play an essential role in endothelial cell recovery following acute radiation injury [32, 33]. In line with these reports, our data identified the TEK receptor for Ang-1 as a dropout in our screen (**Fig. 1D**). We performed GSEA analysis using the Reactome annotation and found that sgRNAs in pathways necessary for endothelial proliferation, including *MAPK-* and *VEGF-*related pathways, were also depleted in our screens and G2/M cell-cycle related terms were found to be dropouts as well, validating the physiological relevance of the results we obtained with our screening approach (**Supplementary Fig. S7**). Interestingly, we identified Cdc-like kinase 2 (CLK2) loss as the most highly enriched knockout across all three types of human endothelium (**Fig. 1D**). When ranking kinases by radiation-specific effects on endothelial proliferation, *CLK2* was among the highest ranked kinases (**Fig. 1D**), while TEK was found within the lowest ranked group.

## CLK2-targeting compounds act as radioprotective therapeutics for endothelium

Damage to the vascular endothelium following treatment with ionizing radiation is associated with adverse patient outcomes via a variety of mechanisms [19–21, 23, 34], and there are currently no therapeutics to prevent these effects. Fortuitously, many small-molecule inhibitors of *CLK2* have already been described [35–37] and investigation of publicly available expression data confirm that *CLK* isoforms are well-expressed in endothelium-rich tissues (**Supplementary Fig. S8A**). To investigate whether pharmacological *CLK2* inhibitors could function as effective radiation countermeasures, we treated HUVEC cells with the small molecule *CLK* inhibitor TG003 2 hours after exposure to 4 Gy radiation [37]. Importantly, TG003 prevented radiation-induced growth inhibition and significantly increased endothelial cell numbers in a dose-dependent fashion (**Fig. 2A; Supplementary Fig. S8B**). Similar effects were observed with Cirtuvivint, which inhibits *CLK2* more selectively than other *CLK* proteins [35] (**Fig. 2B**). These results appeared to be at least partly due to a reduction in ionizing radiation-induced apoptosis as there was a significant decrease in the frequency of endothelial cells positively stained with Annexin V by flow cytometry (**Fig. 2C**) following treatment with TG003, Cirtuvivint, or Lorecivivint, which is another small molecule *CLK2* inhibitor.

**Figure 2.**
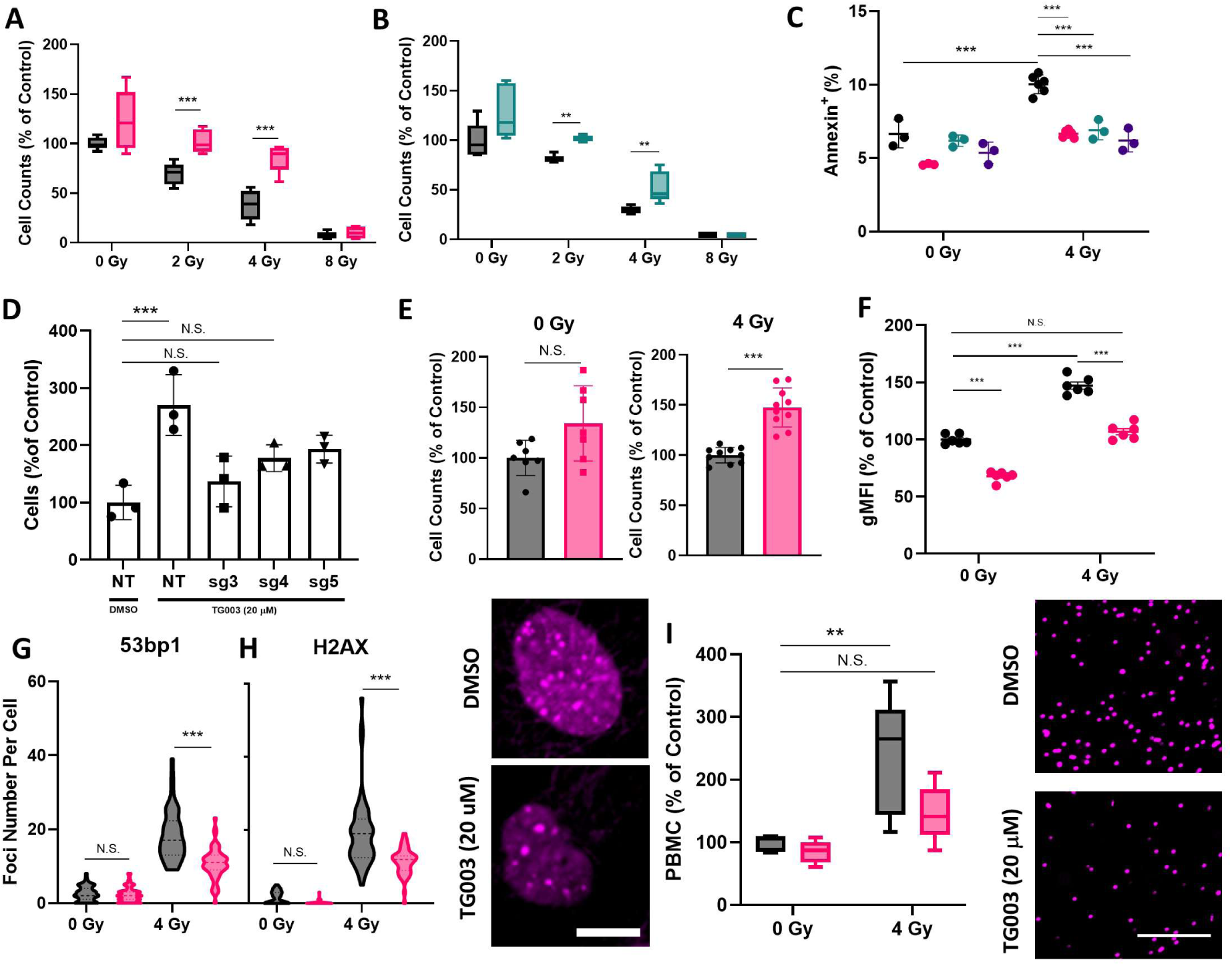
Identification of *CLK2* inhibition as a radioprotective therapeutic strategy. (**A**) Recovery effect of *CLK* inhibition on cell number (cell count as a percentage of control) 7 days post-irradiation when treated with TG003 or (**B**) Cirtuvivint in HUVECs exposed to 0, 2, 4, and 8 Gy of ionizing radiation (n = 6; data are indicative of 2 independent experiments). (**C**) CLK2 inhibitor TG003 significantly reduces percent of Annexin V-positive apoptotic cells measured at 72-hours post-irradiation (n = 3-6 replicates per condition, data are indicative of 2 independent experiments). (**D**) Effect of CLK2 knockout on radioprotective effect of TG003 with 3 different sgRNAs compared to vehicle (DMSO) controls. CRISPR/Cas9-induced loss of CLK2 function abrogates recovery effect of TG003 (NT = non-targeting sgRNA, sg1-3 = CLK2-targeting sgRNA; n = 3). (**E**) Recovery effect of *CLK* inhibition when HSIMECs exposed to 0 or 4 Gy radiation were treated with TG003 (n = 6-9, data are indicative of 2-3 independent experiments). (**F**) Quantification of intracellular reactive oxygen species (ROS) levels by CellROX Deep Red Reagent in HUVEC 24 hours after exposure to 0 or 4 Gy, after treatment with DMSO or TG003 (n = 6 replicates, data are indicative of 2 independent experiments). (**G**) Nuclear fluorescent foci counts for 53bp1 and (**H**) γ-H2AX as well as representative immunofluorescence microscopic images of γ-H2AX foci of HSIMEC exposed to 4 Gy of ionizing radiation and treated with TG003 or DMSO vehicle (bar, 10 μm). (**I**) TG003 reverses leukocyte adhesion induced by 4 Gy of ionizing radiation to endothelial cells 24 hours post-irradiation. ***, p < 0.001; ** p < 0.01; * p < 0.05; N.S., not significant. Colors: Grey/Black = Control, Red = TG003-treated (20 μM), Green = Cirtuvivint-treated (10 nM), Purple = Lorecivivint-treated (100 pM).

Endothelial cells are known to undergo hypertrophy following exposure to ionizing radiation. We similarly observed a radiation dose-dependent increase in cell size, as detected by flow cytometry, which was prevented by TG003 treatment (**Supplementary Fig. S8C**) To confirm that the effects of TG003 were mediated by CLK2 inhibition, we transduced HUVECs with 4 sgRNAs targeting CLK2 or a non-targeting control. Loss of CLK2 via genetic inhibition also promoted cell survival following exposure to ionizing radiation, while treatment with TG003 showed no additional benefit, confirming inhibition of CLK2 activity as the mechanism of action underlying its radiation protecting action (**Fig. 2D)**. Moreover, similar effects were observed in HSIMECs with TG003 increasing cell numbers as compared to DMSO controls following treatment with ionizing radiation (**Fig. 2E**).

As the production of ROS is a key mediator of radiation-induced endothelial cell injury [19], we also explored whether *CLK2* inhibition might promote clearance of ROS. Indeed, staining with CellROX fluorescent dye 24 hours post-irradiation revealed that CLK2 inhibition reduced ROS to untreated levels (**Fig. 2F**). Given the observed reduction in cellular ROS levels, it is possible that *CLK2* inhibition could reverse other downstream consequences of exposure to ionizing radiation. We therefore assessed the formation of γH2AX and 53bp1 foci (two markers of double-stranded DNA breaks) within endothelial cell nuclei, with and without *CLK2* inhibition. Interestingly, *CLK2* inhibition reduced the numbers of both γH2AX and 53bp1 foci per cell, hence suppressing DNA damage in HSIMECs 6 hours post-irradiation (**Fig. 2G-H** and **Supplementary Fig. S9**). Importantly, the reduced levels of ROS and double-stranded breaks produced by treatment with TG003 also resulted in a reduction in endothelial cell-leukocyte adhesion 24 hours post-irradiation (**Fig. 2I**), which should correspond to a reduction in inflammation if similar effects occur *in vivo*.

## CLK2 modulation produces different effects in human endothelial cells versus cancer cells

An important potential clinical application for radiation countermeasure drugs is the mitigation of side effects in cancer patients receiving radiotherapy [38]. While our results with endothelial cells suggest that *CLK2* inhibition could be useful for this application, the opposite response has been observed in cancer cells in one study: *CLK2* overexpression (rather than loss) protected against radiation injury and cell death in cancer-derived HeLa cells[39]. In fact, when we re-analyzed publicly available data from Project Achilles, an atlas of forward genetic screens across the cancer cell line encyclopedia (CCLE) [40], we found that CRISPR knockouts of *CLK2* using the same library we used here also resulted in tumor cell killing (**Supplementary Fig. S10A**). Moreover, analysis of available datasets from the PRISM assay, which assesses the effects of compound treatments on a pool of barcoded cell lines [41], revealed that TG003 treatment either inhibited or had no effect on proliferation of various cancer cell lines. We confirmed these results experimentally using a human tumor-derived Caco-2 intestinal epithelial cell line where we found that TG003 had no protective effect against exposure to ionizing radiation (**Supplementary Fig. S10C**). Thus, CLK inhibition appears to have different and sometimes opposite effects in primary human endothelial cells versus cancer cells.

## CLK2 Inhibition Reverses Radiation-Induced Changes to the Transcriptome and Phospho-proteome

To understand the global effects of *CLK2* inhibition on endothelial cell function following exposure to ionizing radiation, we performed RNA-sequencing of radiation treated and untreated cells, with and without TG003. We identified 923 differentially regulated genes (485 upregulated and 438 downregulated) when we compared cells exposed to 0 versus 4 Gy in the absence of drug, while there were 891 upregulated and 1474 downregulated genes when the TG003 and control (DMSO) treatment groups exposed to 4 Gy ionizing radiation were compared. Many of the most upregulated genes between the 0 Gy and 4 Gy conditions were also downregulated in the 4 Gy TG003 versus DMSO conditions (**Fig. 3A, Supplementary Fig. S11A**). To quantify this relationship, we plotted the effect sizes of all genes comparing 4 Gy TG003 versus DMSO to 0 Gy versus 4 Gy DMSO. We observed a negative correlation (Pearson’s R = −0.555, p < 2.2*10^−16^) in fold changes across all identified transcripts (**Fig. 3A, Supplementary Fig. 11B**), indicating a reversal of the radiation-associated transcriptomic changes. In agreement with this observation, principal component analysis (PCA) of these data showed that irradiated TG003-treated samples associated more closely with control samples that were not exposed to radiation than untreated irradiated samples (**Supplementary Fig. S11D**).

**Figure 3.**
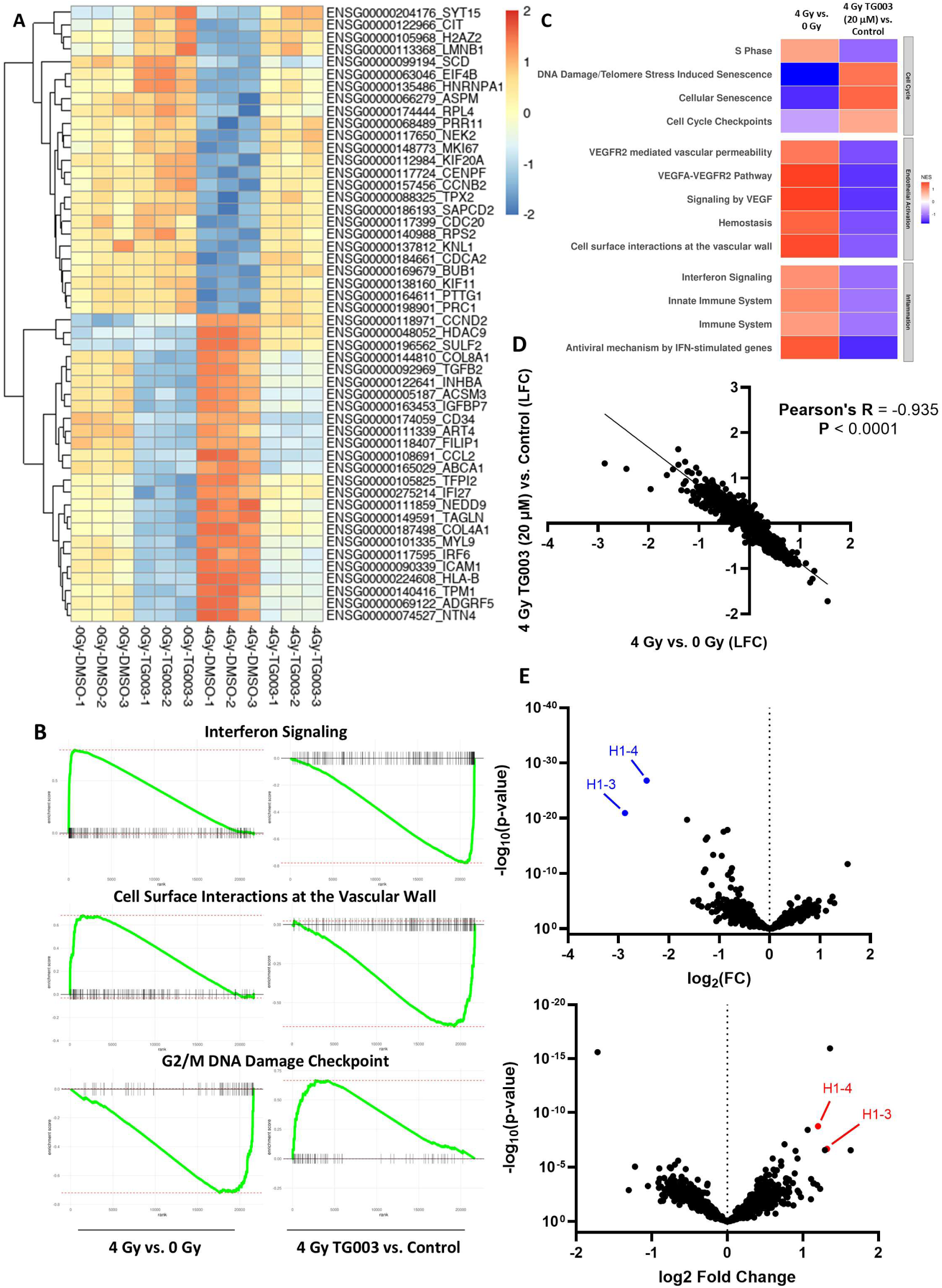
RNA-sequencing and Phospho-Proteomics indicates a reversal of radiation-induced gene signatures. (**A**) Heatmap showing that transcriptional changes induced by 4 Gy of ionizing radiation are reversed by TG003 treatment across all RNA transcripts in human umbilical vein endothelial cells (HUVECs). Heatmap values represent normalized count values. (**B**) Gene set enrichment analysis (GSEA) plots of RNA-sequencing results showing 4 Gy irradiated versus control (left) and TG003-treated and irradiated endothelial cells versus vehicle-treated and irradiated endothelial cells (right) using Reactome annotation (Labels indicate Reactome terms). (**C**) Heatmap (Top) showing significantly altered pathways in GSEA of phospho-proteomic data comparing 4 Gy irradiated versus 0 Gy control HUVECs (left column) and 4 Gy irradiated, TG003-treated versus 4 Gy control HUVECs (right column) 24 hours post-irradiation. (**D**) Scatterplot (bottom) showing log-fold changes of all proteins detected in phospho-proteomic analysis of 4 Gy irradiated versus 0 Gy control HUVECs (X-axis) and 4 Gy irradiated, TG003-treated versus 4 Gy control HUVECs (Y-axis) 24 hours post-irradiation are inversely correlated. (**E**) Volcano plot showing 4 Gy irradiated vs. 0 Gy control (top) and TG003-treated and irradiated endothelial cells versus vehicle-treated and irradiated endothelial cells (bottom). Coloring: Blue = significantly reduced levels, Red = significantly increased levels.

We next sought to understand if these global changes in gene expression were associated with relevant biological responses by performing Gene Set Enrichment Analysis (GSEA) using the Reactome annotation. This analysis revealed a negative normalized enrichment score (NES) for terms relating to Cell Cycle driven by many well-characterized proliferation-related genes (*CCNA2, CCNB2, CCNB1, CDCA8, CDK1*) in irradiated versus non-irradiated cells (**Fig. 3B, Supplementary Fig. S11C,** and **Supplementary Table 4)**. In addition, we observed a positive NES for terms relating to an active inflammatory response (**Fig. 3B, left, Supplementary Fig. S11C, and Supplementary Table 4**) driven by many known inflammatory mediators (*IRF9, IFIT1, HLA-F, OAS2, HLA-B, IFIT2, JAK1, STAT1, STAT2, ISG15*) (**Fig. 3B** and **Supplementary Fig. S11C**), which is consistent with reports that ionizing radiation induces inflammation in endothelial cells [19, 24, 42–45]. We also observed Reactome terms relevant to extracellular matrix organization, in line with radiation acting as a driver of vascular remodeling (**Supplementary Fig. S11C**)[19].

Importantly, when the same analysis was carried out comparing cells treated with *CLK2* inhibitor versus control in the presence of ionizing radiation, we detected a reversal of radiation-associated Reactome terms across the transcriptome (**Fig. 3B** and **Supplementary Fig. S11C**) in agreement with the gene level changes we observed (**Fig. 3B, left** and **Supplementary Fig. S11B**). Indeed, in line with our experimental data modulating *CLK2* activity, we observed reversal of the injury phenotype as indicated by a positive enrichment of Cell Cycle-related terms (**Fig. 3B**). Consistent with our observation that *CLK2* inhibition reduces endothelial-leukocyte adhesion post-irradiation, we also observed down-regulation of terms related to immune/inflammatory pathways and cell surface interactions, in particular *ICAM1* (**Fig. 3A,B**). Intriguingly, a negative NES score was seen in pathways relating to extracellular matrix organization, indicating that *CLK2* inhibition could possibly mitigate this damaging effect of acute radiation exposure as well (**Supplementary Fig. S11C**).

To understand protein-protein interactions that could mediate the radioprotective effect of *CLK2* inhibition, we performed phospho-proteomic analysis of cells treated with and without ionizing radiation, and with and without TG003, 24 hours post-irradiation. In agreement with our RNA-sequencing results, exposure to ionizing radiation resulted in decreased cell cycle-related phosphorylation changes, as well as increased phosphorylation changes associated with an inflammatory response and endothelial cell activation (**Fig. 3C**). In the same manner, TG003 treatment resulted in increased cell-cycle related protein-phosphorylation changes and decreased terms associated with inflammation and endothelial cell activation. These changes were driven by a striking reversal (Pearson’s R: −0.935, p < 0.0001) of proteome-wide changes in protein phosphorylation induced by exposure to ionizing radiation due to treatment with TG003 (**Fig. 3D**). Differences in cell cycle pathways appeared driven at least in part by altered phosphorylation of histones H1-3 and H1-4, which are events preceding mitosis (**Fig. 3E**). In line with these changes, phosphorylation of histones H1-3 and H1-4 was increased by TG003 treatment, contributing to a positive NES score of cell cycle terms. We conclude that TG003 treatment reverses early cellular responses to ionizing radiation across the phospho-proteome, leading to a transcriptome-wide reversal of phenotypes by 7 days. As a profound phospho-proteomic reversal occurs 24 hours post-irradiation, our data suggest that TG003 exerts its protective effects by altering early responses to ionizing radiation. Since we previously observed decreased ROS levels and double stranded break-induced H2AX and 53bp1 foci, these results suggest that TG003 may stimulate ROS clearance and DNA break repair via CLK2 inhibition, thus preventing cell cycle arrest and expression of pro-inflammatory genes (**Supplementary Fig. 10D**).

## CLK Inhibition Protects Against Radiation Injury in Human Intestine and Lung Chips

We next sought to understand if our observations with cultured endothelial cells were relevant in a human tissue- and organ-relevant context. Our group and others have previously demonstrated that the degree of radiation injury in the intestinal epithelium is significantly influenced by the adjacent microvascular endothelium when it is also irradiated [10, 13, 21]. To test if *CLK2* inhibition can protect against functional consequences of ionizing radiation exposure in human intestinal epithelium in an organ-relevant context, we utilized our previously described two-channel microfluidic human Intestine Chip that is lined by primary human ileum organoid-derived intestinal epithelial cells that form highly differentiated intestinal villi when interfaced with primary HSIMEC across a porous membrane, cultured under flow, and exposed to cyclic peristalsis-like mechanical deformations on-chip [46](**Fig. 4A**).

**Figure 4:**
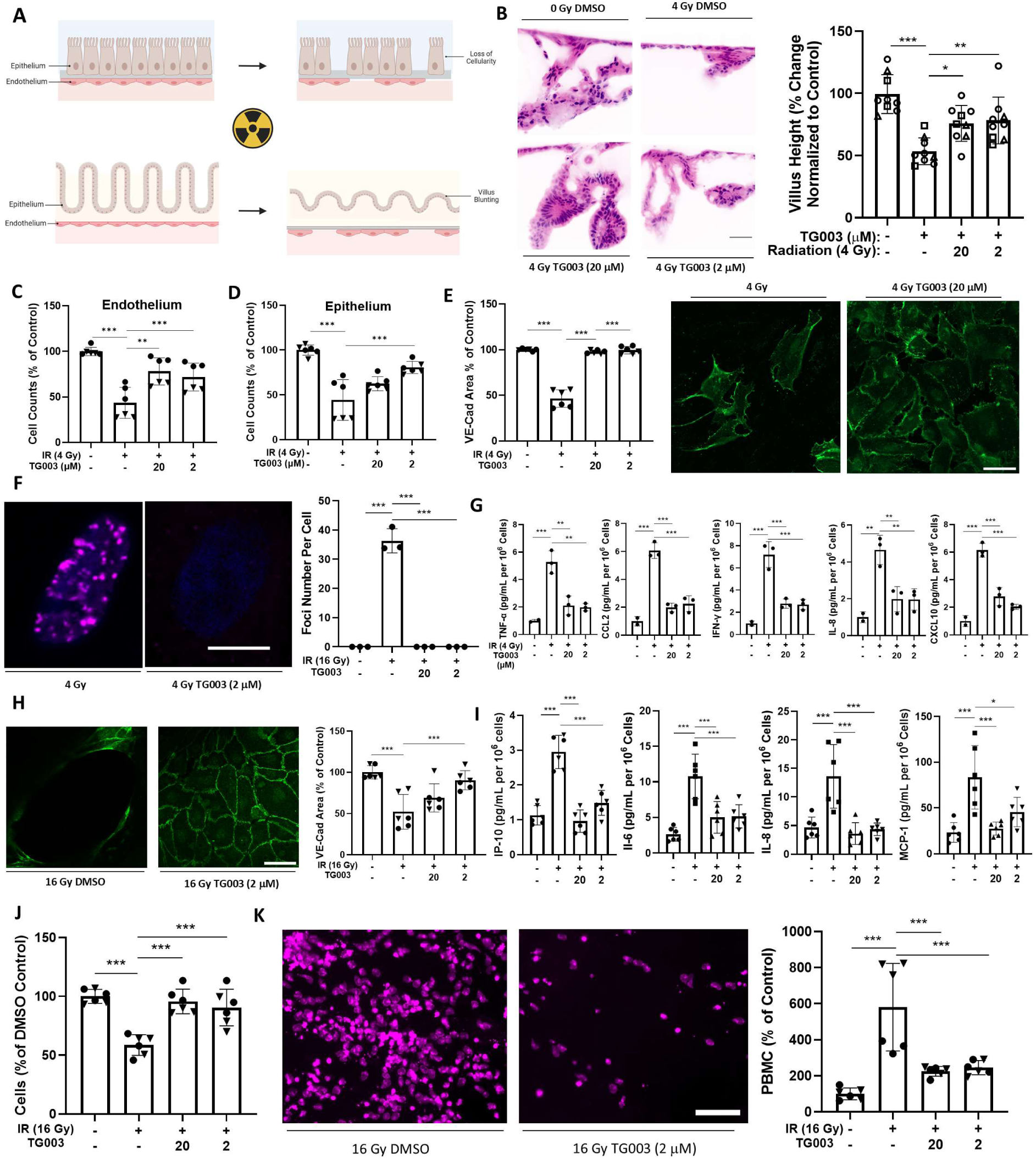
Assessment of CLK2 inhibition as a therapeutic strategy in more physiologically relevant human Intestine and Lung Chips. (**A**) Schematic showing effects of radiation injury in intestine and lung chips. In both systems, exposure to ionizing radiation causes reduced cellularity and loss of tissue architecture. (**B**) Representative pseudo-H&E images (left) and quantification of villus heights (right) across chip cross-sections in intestine chips treated with 4 Gy ionizing radiation or mock and TG003 or DMSO control (n = 9; 3 independent experiments; scale bar, 100 μm; symbols: circle = patient 1, triangle = patient 2, square = patient 3). (**C**) Total cell number counts from the vascular and (**D**) epithelial channels showing TG003 protects against radiation-induced loss of cellularity (n = 6; 2 independent experiments; symbols: circle = patient 1, triangle = patient 2). (**E**) Representative images of VE-Cadherin (left) and quantification of area occupied by VE-Cadherin (right) of control and 4 Gy irradiated intestine chips with vehicle or TG003 7 days post-irradiation (Green = VE-Cadherin; n = 6; 2 independent experiments; symbols: circle = patient 1, triangle = patient 2; scale bar, 50 μm). (**F**) Representative images (left) and quantification (right) showing 53bp1 foci persist in small intestine microvascular endothelial cells treated with 4 Gy ionizing radiation 7 days after irradiation, but not in 4 Gy irradiated, TG003-treated cells (Blue = Hoescht 33342, Magenta = 53bp1; n = 3; 2 independent experiments, scale bar = 10 μm). (**G**) Levels of cytokines detected in effluents of microvascular endothelium channel of human intestine chips exposed to 4 Gy radiation or mock and TG003 or DMSO control by Luminex assay. (n = 3; data are indicative of 2 independent experiments). (**H**) Representative images of VE-Cadherin (left) and quantification of area occupied by VE-Cadherin (right) of control and 16 Gy irradiated lung chips with vehicle or TG003 7 days post-irradiation (Green = VE-Cadherin; n = 6; 2 independent experiments; scale bar, 50 μm; symbols: circle = patient 1, triangle = patient 2). (**I**) Levels of cytokines IP-10, Il-6, Il-8, and MCP-1 detected in effluents of microvascular endothelium channel of human lung chips exposed to 16 Gy radiation or mock and TG003 or DMSO control by Luminex assay. (n = 6; 2 independent experiments; symbols: circle = patient 1, triangle = patient 2). (**J**) Quantification of polymorphonuclear blood cell (PBMC) counts adhered per chip (left) and representative immunofluorescent images (right) of PBMCs adhered to the surface of the microvascular endothelium in human Lung Chips exposed to 16 Gy radiation or mock and treated with TG003 or DMSO controls (Magenta = PBMC whole cell stains; right panel, n = 6; data are indicative of 2 independent experiments; scale bar, 50 μm). (**K**) Cell counts of lung microvascular endothelial cells in chips treated with 16 Gy irradiation or mock and TG003 or DMSO controls, quantified by nuclear staining and imaging. Nuclear counts were assessed in ImageJ (n = 6; 2 independent experiments; scale bar, 200 μm). ***, p < 0.001; ** p < 0.01; * p < 0.05.

When TG003 was flowed through the basal vascular endothelium-lined channel for 7 days beginning 2 h following exposure to 4 Gy ionizing radiation to simulate intravenous administration of this compound in a therapeutic context, we found that *CLK2* inhibition protected against radiation-induced villus blunting in the adjacent intestinal epithelium (**Fig. 4B**). TG003 also reduced both intestinal epithelial cell and endothelial cell loss (**Fig. 4C,D**), which is seen in response to radiation exposure in human intestine *in vivo*. Treatment with ionizing radiation also resulted in disruption of the endothelial monolayer in the Intestine Chip, as observed in capillary beds of patients exposed to ionizing radiation, and TG003 prevented this injury (**Fig. 4E**)[47]. Intriguingly, we observed that intestinal endothelial cells lining the vascular channel of the Intestine Chip and treated with 4 Gy ionizing radiation displayed persistent 53bp1 foci in their nuclei 7 days post-irradiation, which we did not observe in epithelial cells seeded in the same devices that received the same radiation exposure (**Fig. 4F**). However, in chips treated with TG003, we did not observe any 53bp1 foci, reminiscent of unirradiated control cells. Moreover, in accordance with data from our *in vitro* cultures with endothelial cells alone, TG003 treatment reversed radiation-induced increases in pro-inflammatory cytokine levels in the effluents of the vascular channels of the human Intestine Chips (**Fig. 4G**). Thus, *CLK2* inhibition of the endothelial response to radiation exposure can protect human intestinal epithelium from injury in a physiological organ-relevant context.

To understand if this therapeutic strategy may be generalizable to other highly vascularized and radiosensitive tissues, we utilized our previously described human Lung Chip model of radiation injury[11]. Similar to the results on observed in the Intestine Chip, TG003 treatment ameliorated radiation-induced loss of VE-Cadherin expression and junctional continuity within the endothelium in the Lung Chip (**Fig. 4H**). TG003 treatment decreased production of radiation-induced cytokines IP-10, Il-6, Il-8, and MCP-1 in the vascular channel effluents of irradiated Lung Chips as well, similar to our results from the Intestine chip (**Fig. 4I**). Protection against radiation injury was also evidenced by maintenance of increased numbers of viable pulmonary vascular endothelial cells (**Fig. 4J**). In line with this result, TG003 treatment reduced adhesion of circulating peripheral blood mononuclear cells (PBMCs) that were perfused through the vascular endothelium following exposure to ionizing radiation (**Fig. 4K**). Thus, *CLK2* inhibition protected against radiation injury in at least two different human organ models.

## Discussion

Taken together, these findings indicate that pharmacologic CLK2 inhibitors potentially could be repurposed as radioprotectants in multiple tissues and organs and hence be useful for treatment of patients with ARS. Our results also demonstrate the ability of using CRISPR screens to identify molecular targets that mediate human vascular endothelial cell-specific sensitivity to radiation injury as well as to rapidly identify existing drugs that may repurposed as radiation countermeasures, which can then be validated experimentally using human Organ Chips. This was accomplished using cell survival and continued proliferation as readouts in primary human endothelial cells isolated from multiple organ sources. However, cell growth represents only one biological response to ionizing radiation exposure, and thus, this same approach could be used to screen for alternative readouts (e.g., formation of ROS, radiation-associated inflammation, extracellular matrix alterations) in endothelial cells or other radio-sensitive cell types in future studies.

The cellular functions of CLK2 remain poorly understood. While one past study showed that CLK overexpression promotes cancer-derived HeLa cell survival and growth after treatment with ionizing radiation [39] and others showed that CLK2 and CLK4 inhibition inhibit tumor-derived Caco-2 cell proliferation [48], we observed no effect of CLK2 inhibition of Caco-2 cells and that it sustained, rather than inhibited, cell growth in radiation-exposed endothelial cells. A possible explanation for this discrepancy is that we treated Caco-2 cells with only 4 Gy of ionizing radiation to see a synergistic effect preventing cell survival, whereas the previous study treated cells with 10 Gy. Nonetheless, our data clearly demonstrate that CLK2 inhibition promotes survival and growth in multiple types of healthy endothelial cells exposed to radiation, whereas it appears to have either minimal effects or promote death in various cancer cell lines. Importantly, this could have great clinical significance as it suggests that CLK2 inhibitors might offer a novel combined anti-cancer and radioprotective therapeutic strategy by inducing tumor cell killing while protecting healthy cells against radiation injury in cancer patients receiving concurrent radiotherapy.

In addition to its effects on cell survival and growth, inhibition of CLK2 induced transcriptome- and phospho-proteome-wide changes in signaling, as well as a decrease in radiation-induced leukocyte adhesion to the vascular endothelium. These data are highly encouraging but were acquired by studying one cell type at a time in a conventional static culture model. Therefore, it was necessary to reproduce these results in a more physiologically relevant system. For this reason, we tested CLK2 inhibition in microfluidic human Intestine and Lung Chip models of acute radiation injury, which we have previously shown recapitulate many features of radiation induced injury observed in humans in vivo, including organ-specific radiation dose sensitivities [10, 11]. Intriguingly, we also observed the presence of persistent 53bp1 foci in endothelial cells, but not epithelial cells exposed to ionizing radiation, which was abrogated completely by treatment with TG003. The presence of these persistent foci could represent an endothelial-specific mechanism of radiation injury, which TG003 prevents. The finding that treatment with CLK2 inhibitors reduced radiation injury in both Organ Chips validated our earlier results with cultured endothelial cells alone. By testing in human Organ Chips, we also were able to perform our entire drug repurposing effort rapidly, from target identification to therapeutic testing in a fully human context. While testing in animal models enables observation of responses in a whole organism, these benefits are offset by animal-to-human differences in drug toxicities and efficacies[49]. For this reason, the FDA will now consider data generated in human relevant in vitro models, including human Organ Chips, in future investigational new drug (IND) applications in lieu of data from animal studies[50, 51]. Thus, these findings may help to accelerate the delivery to the clinic of an entirely new class of radiation countermeasure drugs that target endothelium found in many radiosensitive organs.

## Author contributions

Conceptualization: RRP, DBC, DEI

Methodology: RRP, AO, YM, JJ, JDL, NTL, JFF

Investigation: RRP, JFF, AO, YM, JDL, AJ, JJ, BB

Visualization: RRP, AMH, CK, JDL

Funding acquisition: DEI, DBC

Supervision: DEI, DBC, BB

Writing – original draft: RRP

Writing – review & editing: RRP, DBC, DEI, CK, AMH

## Data and materials availability

All data are available in the main text or the supplementary materials.

