## Supplementary Figures for "Endothelial CLK2 as a therapeutic target for acute radiation syndrome"

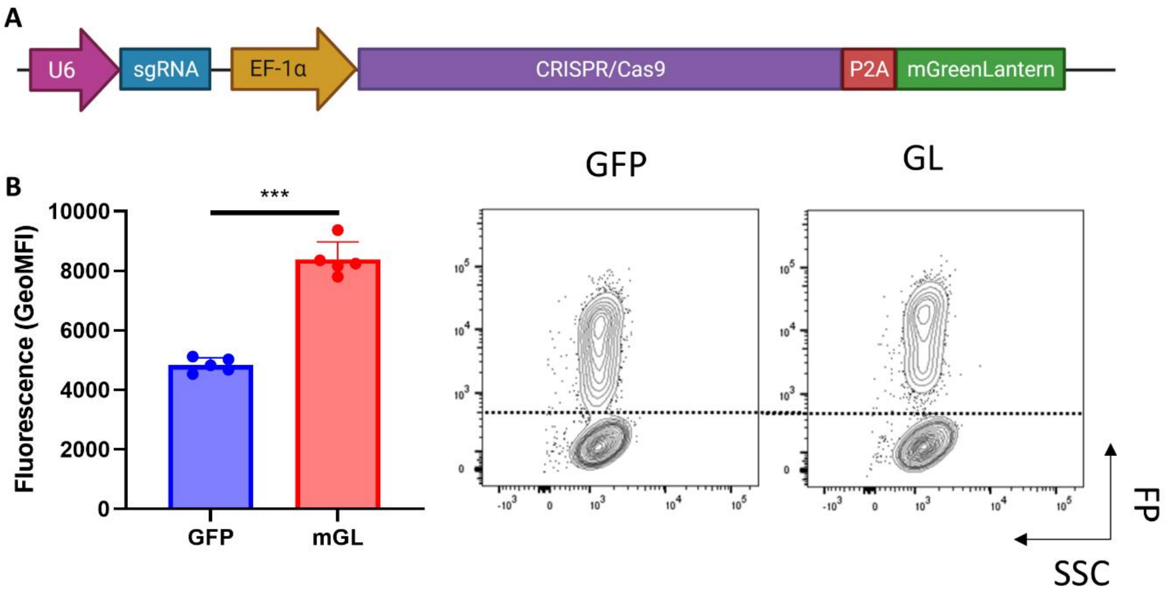


**Supplementary Figure S1: Improved Signal-to-Noise with LCv2-mGL and optimization of a kinome-scale radiation resistance screen in human endothelial cells.** (**A**) Schematic showing key components of LentiCRISPRv2-mGreenLantern. (B) LentiCRISPRv2-mGreenLantern shows superior signal to noise by Geometric Mean Fluorescence Intensity (GeoMFI) as compared to LentiCRISPRv2-GFP assessed by flow cytometry 72 hours post-transduction in K562 cells (n = 5).

**Supplementary Figure 2: Highly Efficient Lentiviral Transduction of Primary Endothelial Cells with LCGL**

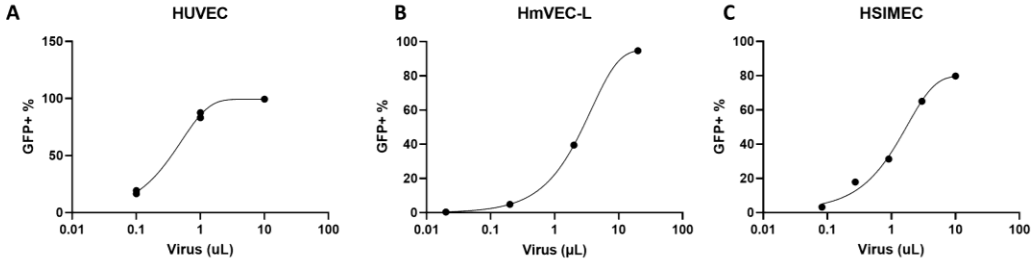


**Supplementary Figure S2: Highly Efficient Lentiviral Transduction of Primary Endothelial Cells with LCGL.** (**A**) Dose titration of concentrated lentiviruses in human umbilical vein endothelial cells (HUVEC), (**B**) human microvascular lung endothelial cells (HmVEC-L), and (**C**) human small intestinal microvascular endothelial cells (HSIMEC).

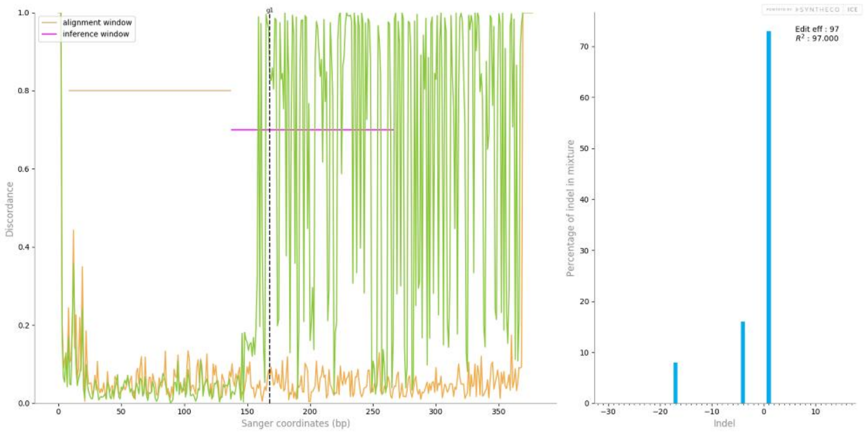


**Supplementary Figure S3: Efficient editing at 7 days post-transduction with LCGL in HSIMEC.** (Left) Plot showing editing efficiency in human small intestinal microvascular endothelial cells transduced with LCGL containing an sgRNA targeting *TP53* as compared to untransduced controls and (Right) estimated editing efficiency by ICE analysis.

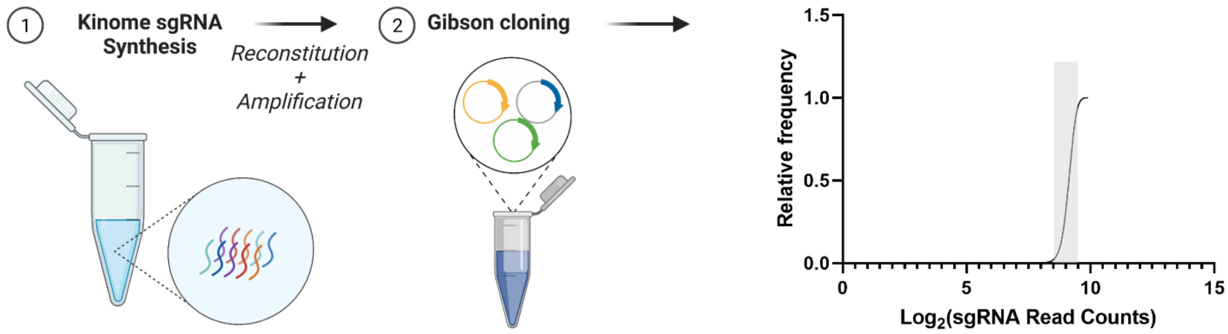


**Supplementary Figure S4: Cloning of Kinome-scale sgRNA Library.** Schematic showing cloning workflow and sgRNA frequency counts. Grey shading indicates sgRNAs within the 90$^{th}$ and 10$^{th}$ percentile.

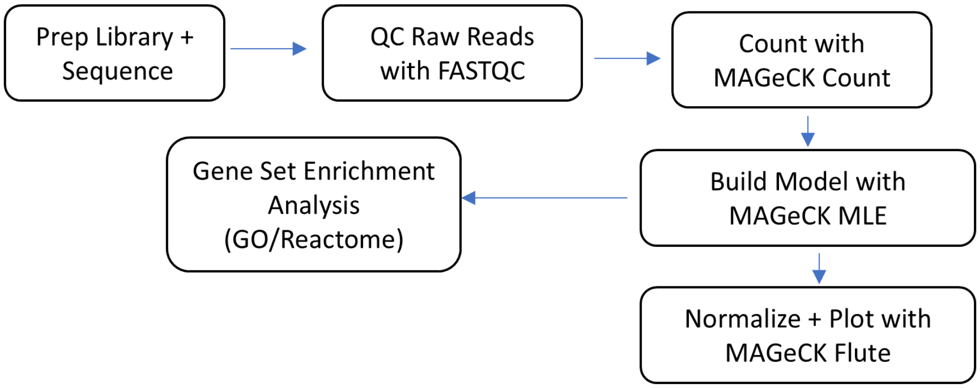


**Supplementary Figure 5: Overview of Screen Analysis Pipeline.** Schematic showing overview of computational analysis pipeline of screen. Raw reads are quality controlled (QC) with FASTQC, raw counts are assessed in MAGeCK Count, and either used for gene set enrichment analysis (GSEA) or normalization and ranking is performed using MAGeCK Flute. All analyses are performed using Galaxy.

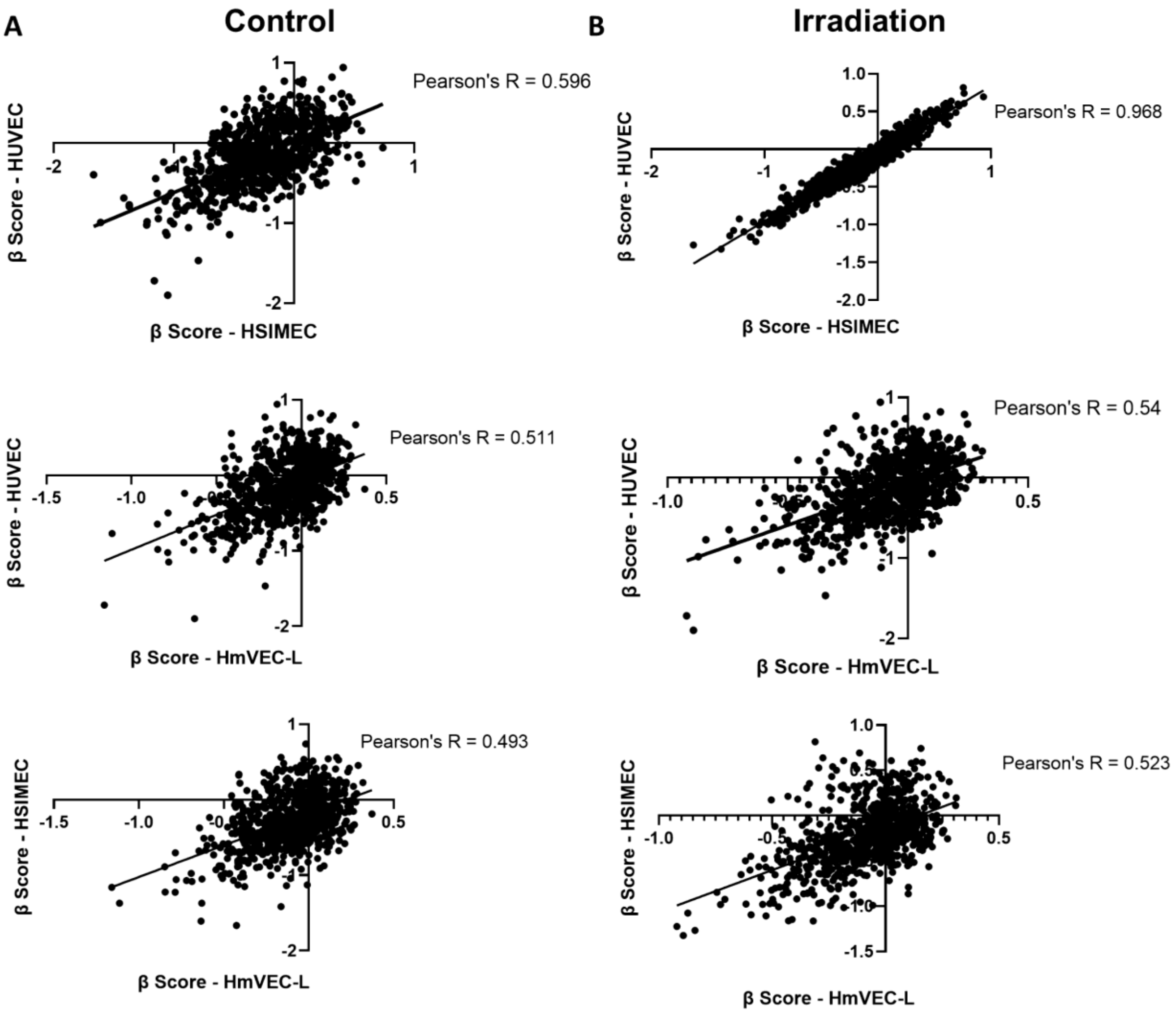


**Supplementary Figure 6: Correlation between results from endothelial cells of differing origin.** Correlation plots showing correlation between β scores of endothelial cells isolated from different tissues of origin under Control (**A**) or Irradiated (**B**) conditions.

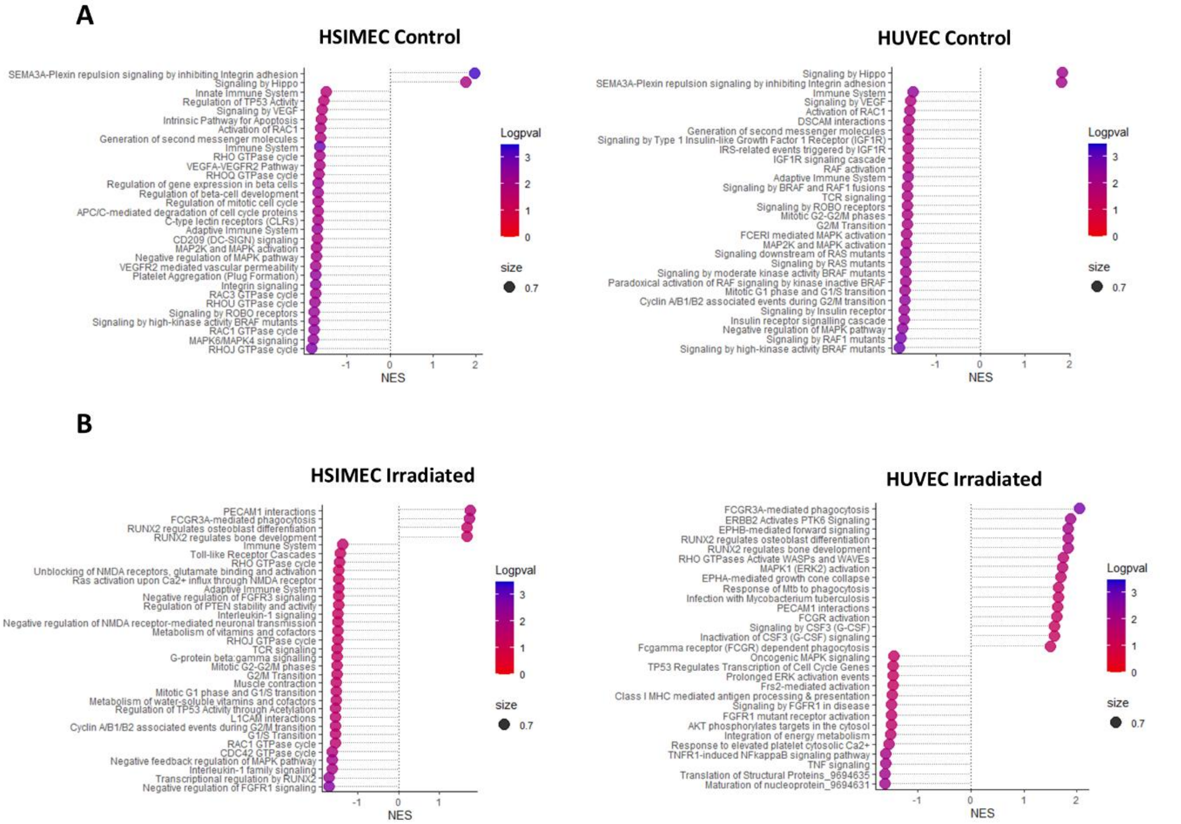


**Supplementary Figure 7: GSEA output of pathway analysis of screen results.** Pathways identified via ranked gene-level analysis of sgRNAs using Reactome annotation under control (**A**) or Irradiated (**B**) conditions. All analyses are performed in Galaxy EU as shown in Supplementary Fig. 5.

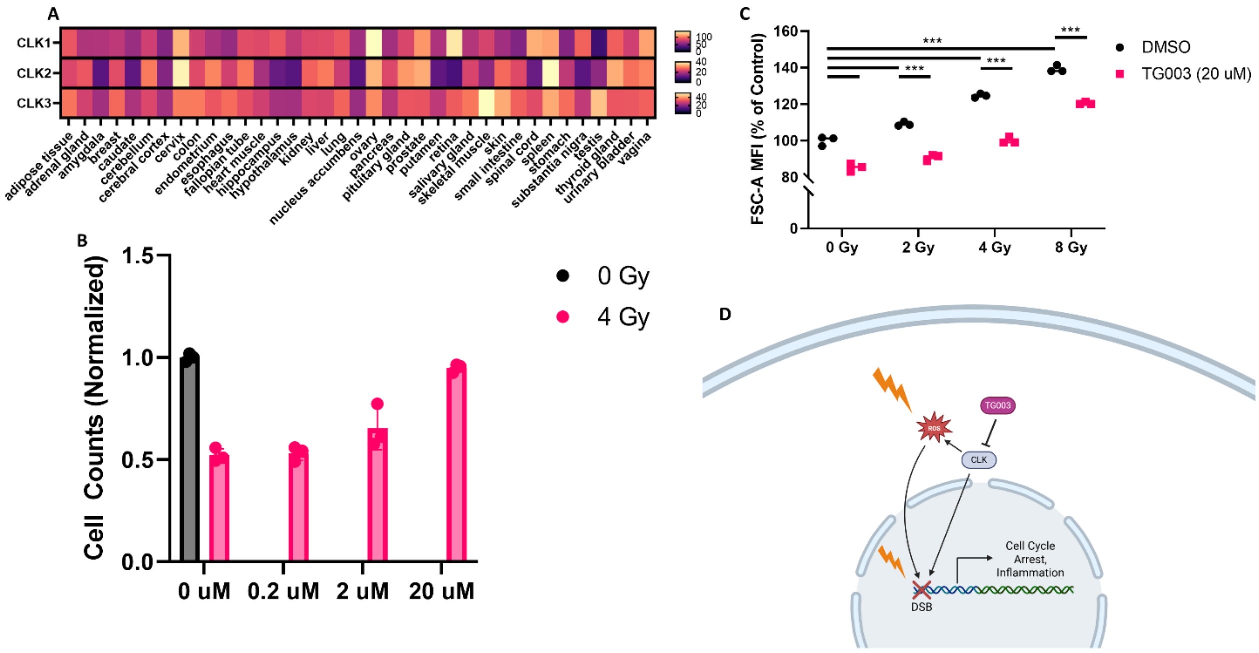


**Supplementary Figure 8: CLK2 is widely expressed in endothelial-rich tissues and modulation affects irradiated endothelial cells**. **(A)** Expression data from GTEx showing high levels of CLK gene expression across human tissues, particularly in vascular-rich tissues. Shading indicates transcript per million counts (TPM). (**B**) Dose response of TG003 in irradiated HUVECs treated with TG003 2 hours post-irradiation and counted at 7 days post-irradiation (n = 3, data are indicative of 2 independent experiments). (**C**) TG003 treatment reduces cell size increases caused by treatment with 4 Gy ionizing radiation. (**D**) Schematic showing proposed model of CLK2 inhibition via TG003, which affects both intracellular ROS levels and 53bp1 and H2AX foci counts.

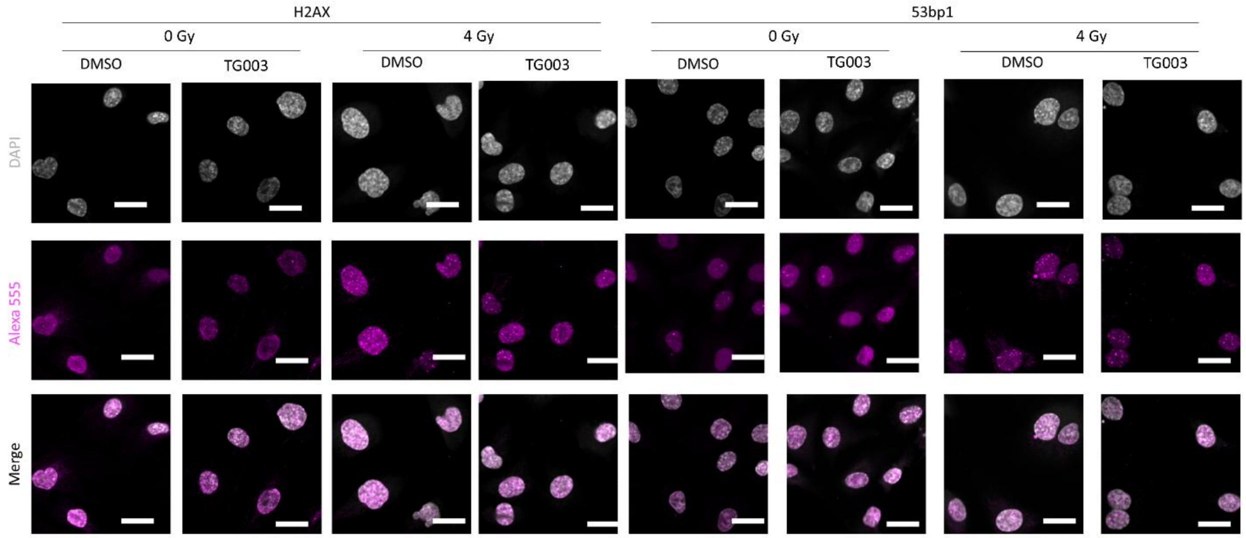


**Supplementary Figure 9: gH2AX and 53bp1 foci are reduced in treated samples vs. controls.** Representative images of nuclei stained for H2AX or 53bp1 using an Alexa 555-conjugated antibody.

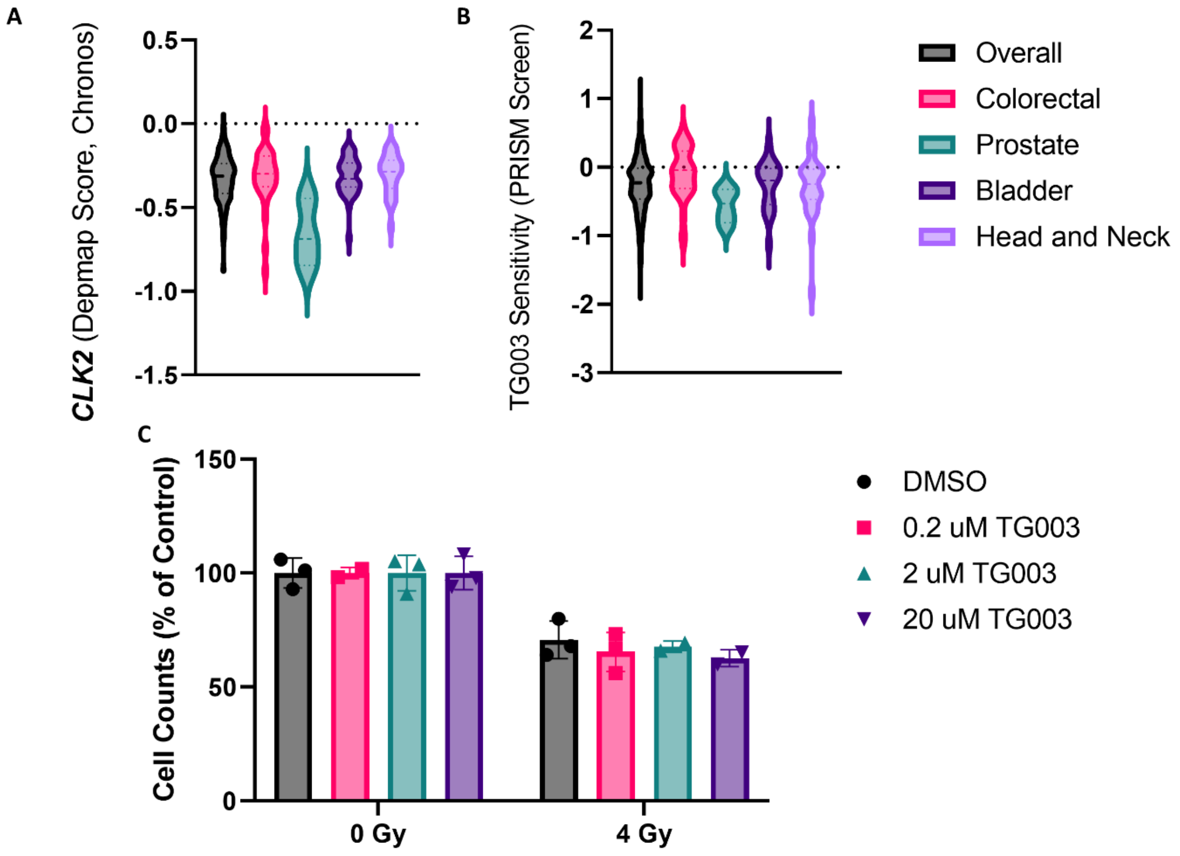


**Supplementary Figure 10: Depmap and PRISM Assay Scores for CLK2 ablation in Cell Lines.** (**A**) Depmap scores for *CLK2* knockout across human cell lines, separated by subtype. Negative scores indicate depleted sgRNAs targeting *CLK2*. (**B**) Effect of TG003 treatment in the PRISM assay. Negative scores indicate depletion of cell lines. (**C**) No additional effect seen in Caco-2 cells treated with TG003 for 7 days after exposure to 4 Gy of ionizing radiation.

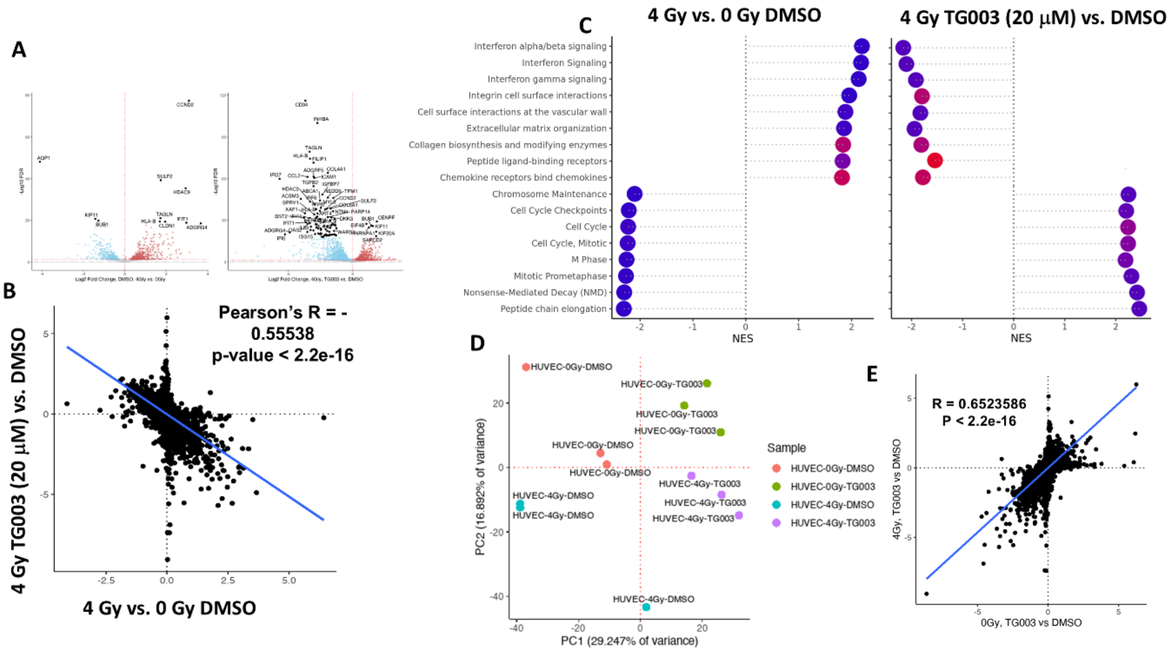


**Supplementary Figure 11: Extended Analysis of RNA-sequencing and phospho-proteomic Data**. (**A**) Volcano plot of 4 Gy vs. 0 Gy treated HUVEC RNA sequencing results (left) and 4 Gy TG003-treated vs. 0 Gy control HUVEC RNA sequencing results (right) 7 days post-irradiation (n = 3). (**B**) Negative correlation between log-fold changes observed comparing 4 Gy vs. 0 Gy DMSO-treated HUVEC (X-axis) and 4 Gy, TG003-treated vs. 4 Gy DMSO-treated HUVEC (Y-Axis) 7 days post-irradiation. (**C**) Significantly enriched Reactome pathways observed in HUVECs treated with 4 Gy ionizing radiation vs. control or 4 Gy ionizing radiation and TG003 vs. DMSO controls. (**D**) PCA plot of phospho-proteomic data showing clustering of 4 Gy TG003-treated samples with 0 Gy control samples as compared to 4 Gy-treated control samples. (**E**) Positive correlation between 4 Gy TG003-treated vs. DMSO control samples (Y-axis) and 0 Gy TG003-treated vs. DMSO control samples (X-axis).

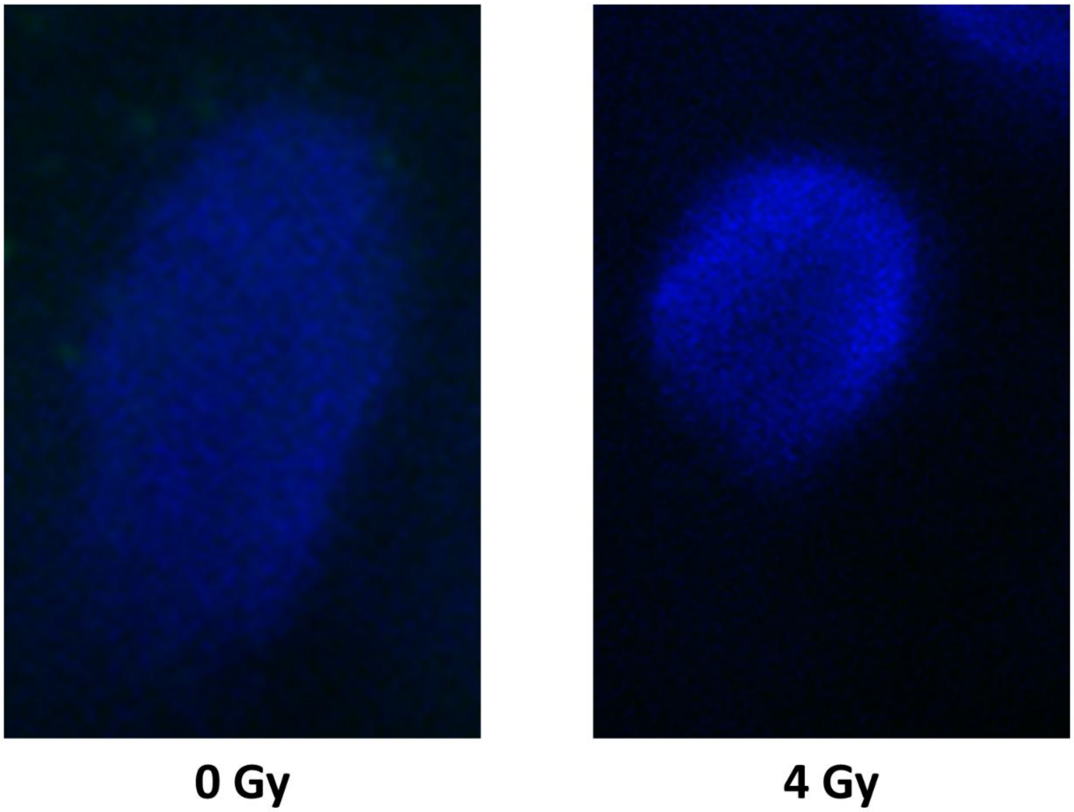


**Supplementary Figure 12: 53bp1 Staining in Intestinal Epithelial Cells 7 days post-irradiation**. Intestinal epithelial cells cultured in human organ chips stained with DAPI (shown in blue) co-stained with 53bp1 (shown in green), show minimal 53bp1 foci. Images were concurrently taken with endothelial cell foci (main Fig. 4F).
